# Development of ENTV reverse genetics system and phenotypic evaluation of rescued virus reveals host-specific replication patterns in mosquitoes

**DOI:** 10.1101/2025.07.29.667424

**Authors:** Marina Fujii, Emily N Gallichotte, Irma Sanchez-Vargas, Brooke M Enney, Lauren E Malsick, Gregory D Ebel, Brian J Geiss

**Affiliations:** Department of Microbiology, Immunology, and Pathology, Colorado State University, Fort Collins, USA; School of Biomedical and Chemical Engineering, Colorado State University, Fort Collins, USA

## Abstract

Entebbe bat virus (ENTV) is a bat-associated flavivirus with no known vector. Research into the biology of this virus, including assessment of the possibility that it may be vector-transmitted, is hindered by a lack of molecular tools and robust genetic systems. Therefore, we sequenced the complete 3’ untranslated region, which was not previously available, and developed an infectious clone of ENTV to facilitate further investigation of the virus. Virus derived from the clone replicated similarly to the parental virus isolate in various vertebrate cells. Surprisingly, ENTV replicated to high titers in *Aedes aegypti* and *Aedes albopictus* mosquito cell lines, but there was no replication or infection in *Culex tarsalis* cells. In addition, phylogenetic and bioinformatics analyses strongly suggested that ENTV may be associated with a mosquito host. Given the bioinformatics support and efficient growth in *Aedes* cells, we orally exposed *Ae. aegypti* and *Ae. albopictus* to ENTV to evaluate infection. The ENTV blood-fed mosquitoes were all negative for infection; however, when ENTV was intrathoracically inoculated, bypassing the initial midgut infection and escape barriers, it replicated to high levels in the body, without dissemination of infectious virus into the saliva. These findings suggest that, despite demonstrating high molecular compatibility at the cellular level in *Aedes* mosquitoes, *Ae. aegypti* and *Ae. albopictus* are unlikely to serve as competent vectors for ENTV transmission due to strong midgut infection barriers. The clone presented in this manuscript should help to clarify the mechanisms for transmission and maintenance of ENTV, which remain poorly understood.

## Introduction

Entebbe bat virus (ENTV) was first isolated from the salivary gland of a little free-tailed bat (*Mops pumilus*) in Uganda in 1957.^1^ ENTV belongs to the genus Orthoflavivirus, which contains medically important pathogens such as dengue virus, Japanese encephalitis virus, and West Nile virus. Orthoflaviviruses can be classified into four groups based on their host specificity: mosquito-borne, tick-borne, insect-specific, and no-known vector (NKV). ENTV is classified as a NKV flavivirus: ENTV has only been isolated twice,^1,2^ both from *Mops pumilus*, despite robust virus surveillance efforts from mosquitoes in the same region.^3–5^

For an arthropod to be a competent vector of a virus, it must be able to become infected after feeding on a viremic animal and then, after an extrinsic incubation period, transmit the virus to a naïve host. Multiple studies have sought to determine if mosquitoes are competent vectors of ENTV.^6,7^ To determine if mosquito cells are susceptible to ENTV, Varelas-Wesley and Calisher (1982) inoculated ENTV on C6/36 (*Aedes albopictus*) cells. Extracellular virus titer was low (∼10^2^ pfu/mL), approximately 5 logs lower than St. Louis encephalitis virus, a known mosquito-borne flavivirus (MBF), concluding inefficient replication of ENTV in these cells.^7^ Simpson and O’Sullivan (1968) blood-fed *Ae. aegypti* mosquitoes with ENTV and tested transmission to mice.^6^ They found infectious ENTV in a pool of whole mosquitoes at 21 days post-infection (dpi)—however, none of the mosquito bites transmitted the virus to mice. Together, these results suggest that while ENTV may infect and replicate in *Aedes* cells and mosquitoes at low levels, *Aedes* mosquitoes are poor/incompetent vectors for ENTV.

Although field and laboratory findings do not show evidence of vector transmission, ENTV has multiple features suggesting the role of an arthropod vector host. In previous analyses using available ENTV sequences (not containing the entire 3’ UTR), ENTV and genetically similar Sokoluk virus (SOKV), and Yokose virus (YOKV) comprise a subclade within the MBF clade, in contrast to other NKV flaviviruses such as Rio Bravo virus (RBV) or Apoi virus, which form a distinct clade branching from a common ancestor with tick-borne flaviviruses.^2,8,9^ In addition, the CpG/UpA dinucleotide usage of ENTV, SOKV, and YOKV—which is known to differ between viruses infecting vertebrates and invertebrates—was comparable to that of arthropod-borne flaviviruses. In contrast, other NKV flaviviruses in the RBV clade exhibited a dinucleotide bias favoring vertebrate hosts, further suggesting that viruses in the ENTV clade may have an arthropod host.^8^

Reverse genetics–generation of infectious virus from cDNA–is a powerful tool in virology, and others have found viral determinants of host-specificity using this approach.^10–12^ Reverse genetics systems have been established for many flaviviruses and are widely used to study the function of viral proteins, generate reporter viruses, and develop chimeric viruses for vaccine development.^13–19^ Flaviviruses have an ∼11kb positive-strand RNA genome with a single open reading frame that encodes a single polyprotein, which is then cleaved into individual proteins post-translationally by viral and host proteases. This genetic design allows one to generate infectious virus by simply introducing viral genomic RNA or cDNA into cells. For RNA *in vitro* transcription, bacterial T7 polymerase is a popular choice. Although the lack of proofreading function in T7 polymerase may contribute to the generation of mutant swarm akin to the field-isolate viruses, transcription and handling of RNAs in the tube require extra reagents and time compared to a simple cDNA transfection that would produce genomic RNAs in transfected cells.

One difficulty complicating this otherwise straightforward method is flavivirus genome instability in bacteria, presumably due to the unexpected protein expression from cryptic promoters.^18^ This issue can often be circumvented by cloning cDNA in separate plasmids/fragments.^13,16,17^ In that case, the partial genome fragments are ligated in vitro followed by RNA transcription^13,14,16^ or cDNA amplification by circular polymerase extension reaction (CPER).^17^ Others succeeded by using a special strain of *E coli*,^14,20,21^ low-copy number plasmids,^14,22^ or by modifying the sequence to disrupt potential cryptic prokaryotic promoters.^18^ Whatever the approach, the resultant reverse genetics system should generate a virus that closely resembles the wildtype while being easy to use.

In this study, we aim to establish an easy-to-use reverse genetics system of ENTV as a tool to further study this phylogenetically unique NKV flavivirus. Furthermore, we examine the possibility that ENTV is transmitted by mosquitoes, which early lab experiments did not show evidence for. In this study, we revisited the mosquito infection experiments with today’s standard methodologies.

## Results

### Infectious clone was recovered efficiently

Because the sequence for ENTV UG125 (KP233893.1) did not contain the complete 3’ untranslated region (UTR), we performed 3’ rapid amplification of cDNA ends (RACE) and determined the sequence of an additional 152nt. RNA secondary structure prediction confirmed the presence of the conserved 3’ stem-loop structure found in 3’ termini of flavivirus UTRs, indicating the coverage of the entire 3’ UTR (**Supplementary Figure 1**). We initially attempted to clone the entire ENTV genome into a single plasmid. However, the product was unstable in *E. coli*. To disrupt the potential cryptic promoters, we cloned the ENTV genome into four plasmids with ENTV amplicons possessing overlapping ends (pBG878, pBG879, pBG880, pBG881). The plasmid containing the ENTV 5’ UTR (pBG878) possesses a cytomegalovirus (CMV) Pol II promoter with the transcriptional start site positioned at the first viral nucleotide. The plasmid containing the 3’ stem loop structure (pBG881) possesses a 3’ hepatitis delta virus ribozyme that cleaves the Pol II transcript immediately after the last viral nucleotide, generating an authentic 3’ end. To generate recombinant ENTV, PCR amplicons were generated from each plasmid, Gibson-assembled, and transfected into BHK cells (**Figure 1A**). Cytopathic effect was observed at 3 days post-transfection (**Figure 1B**), and infectious virus was consistently recovered from three independent transfections despite relatively low transfection efficiency (<10%) measured from the control GFP expression plasmid. In all three independent transfection replicates, the virus titer peaked at 4 days post-transfection, reaching titers greater than 10□ pfu/mL (**Figure 1C**).

**Figure 1.**
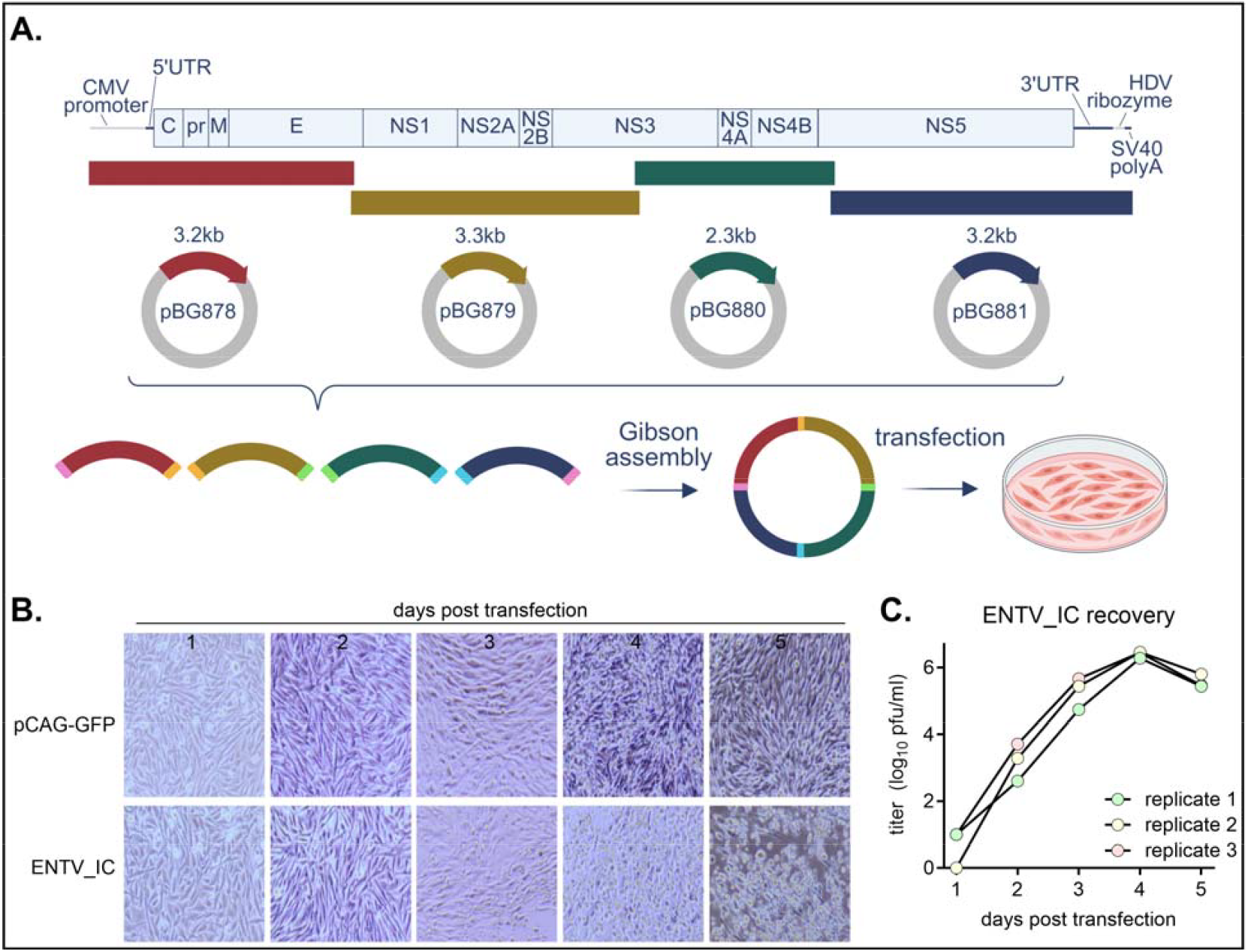
Establishment of ENTV infectious clone. **A)** A schematic image of the reverse genetics system. ENTV genome was split into four fragments and cloned into a CMV promoter-HDV ribozyme cassette. P0 virus was generated by transfecting Gibson assembly product directly to BHK cells. **B)** 500ng of mock plasmid (pCAG-GFP) or ENTV_IC was transfected to BHK cells, and pictures of CPE taken every 24 hours. **C)** Recovery of ENTV_IC from three independent transfections. Cell culture media was collected every 24 hours, frozen at -80°C, and titrated on BHK cells.

### ENTV and ENTV_IC replicated similarly in various cell lines

We first compared virus plaques in BHK cells and found that infectious clone-derived virus (ENTV_IC) had a plaque size comparable to the original ENTV isolate (herein referred to as ENTV) (**Figure 2A, 2B**). The ENTV_IC plaque size and morphology were more homogenous than ENTV, a phenotype consistent with genetically monoclonal infectious clones as opposed to isolates rich in genetic diversity (**Figure 2B**). We next compared virus growth in various mammalian, avian, mosquito, and tick cell lines known to support the propagation of at least one flavivirus. ENTV_IC had comparable replication kinetics to ENTV in all cell lines tested, generating similar levels of extracellular infectious virus. Consistent with previous reports, both ENTV replicated well in Vero E6 (*Chlorocebus sabaeus*)^7^ and BHK (*Mesocricetus auratus*)^2^ cells (**Figure 2C**). ENTV also replicated in many vertebrate cell lines, including AJ6 (*Artibeus jamaicensis*), Huh7 (*Homo sapiens*), and DF-1 (*Gallus gallus domesticus*) (**Figure 2C**). To our surprise, ENTV replicated in multiple *Aedes* cell lines, with peak titers reaching ∼10□ and ∼10□ pfu/mL in Aag2 (*Ae. aegypti*) and C6/36 (*Ae. albopictus*) cells, respectively (**Figure 2D**). In contrast, ENTV failed to produce detectable infectious virus in CT (*Culex tarsalis*) and ISE6 (*Ixodes scapularis*) cells (**Figure 2D**). We confirmed that CT and ISE6 cells can support the replication of relevant viruses using West Nile virus and Powassan virus as positive controls, respectively (**Supplementary Figure 2A**).

**Figure 2.**
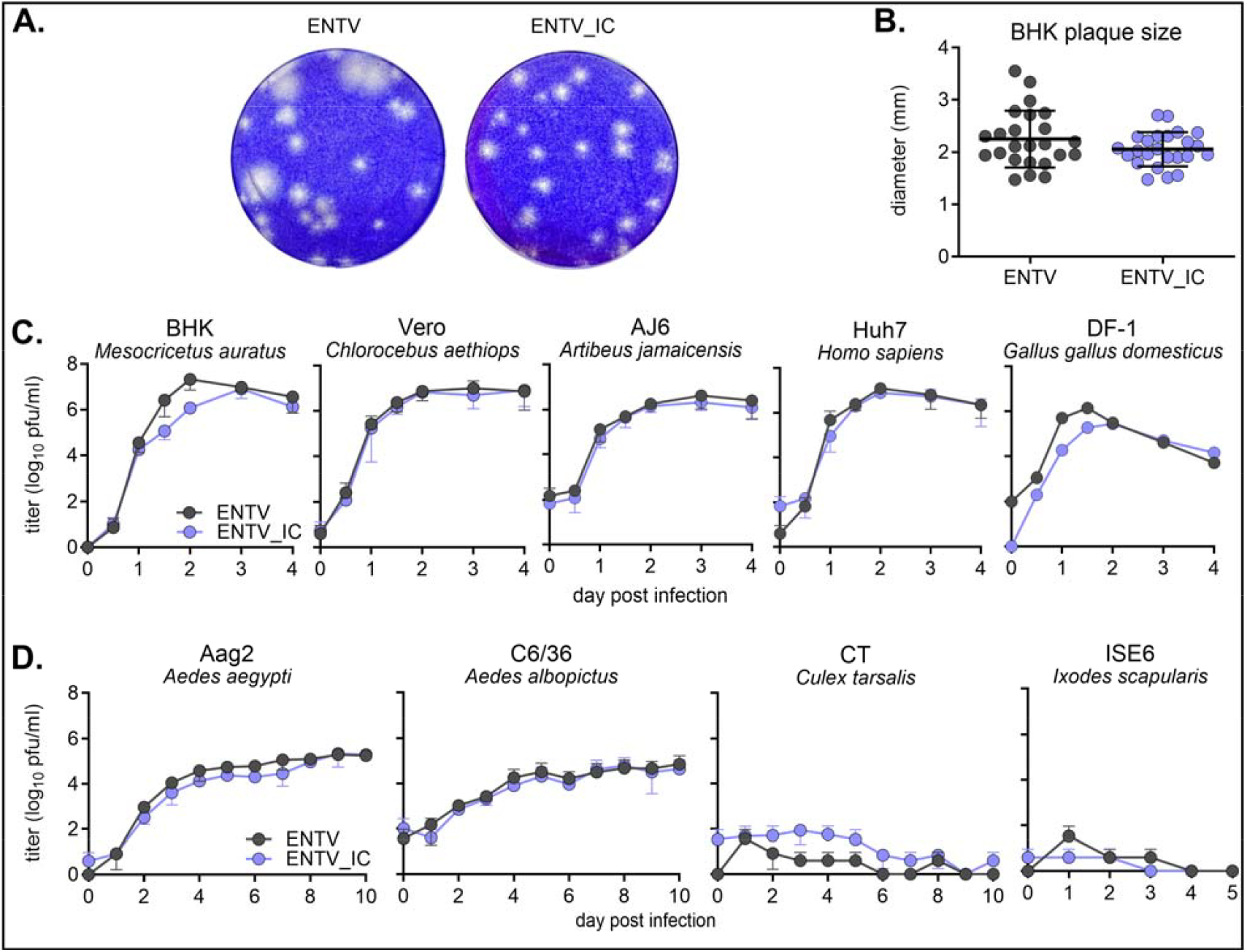
ENTV_IC shows a comparable phenotype as ENTV. **A)** 5dpi plaques on BHK cells. **B)** Plaque diameters of ENTV and ENTV_IC (mean ± SD). No statistically significant difference was detected using Mann-Whitney’s rank test (n=24, p>0.05). **C, D)** Sub-confluent vertebrate (**C**) or invertebrate (**D**) cell monolayers were inoculated with ENTV or ENTV_IC at an MOI of 0.01 in biological triplicate (mean ± SD). Cell culture supernatant was collected at each time point, frozen at -80°C, and titrated on BHK cells.

### ENTV genome characteristics suggest a mosquito host

Because ENTV replicated efficiently in mosquito cells, we assessed whether ENTV likely has a mosquito host through genome sequence analysis. Consistent with previous literature,^2,8,9^ using the complete ENTV containing the new 3’ UTR sequence, ENTV, SOKV, and YOKV form a subclade within MBFs (**Figure 3A**). In the 3′ UTR, MBFs contain two conserved sequences (CS), CS1 and CS2. Having determined the complete 3′ UTR, including a previously uncharacterized region containing the putative CS1, we compared the ENTV sequence with those of other flaviviruses. Consistent with MBF viruses, ENTV had CS1 and CS2, in contrast to NKV flaviviruses in the RBV clade, which do not possess clear CS1 and CS2 (**Figure 3B**). Next, we used a bioinformatics tool (https://bioinformatics.cvr.ac.uk/software/viral-host-predictor/) to computationally predict the host/vector of ENTV. The algorithm considers phylogeny and sequence biases, including codon usage, codon-pair score, and dinucleotide frequencies^23^. The reservoir vertebrate host was predicted to be Vespertilioniformes, a suborder of Chiroptera, including the only known host of ENTV, *Mops pumilus* (**Figure 3C**). The model predicted ENTV as arthropod-borne, with strong computational support for mosquitoes as the most likely vector (**Figure 3C**).

**Figure 3.**
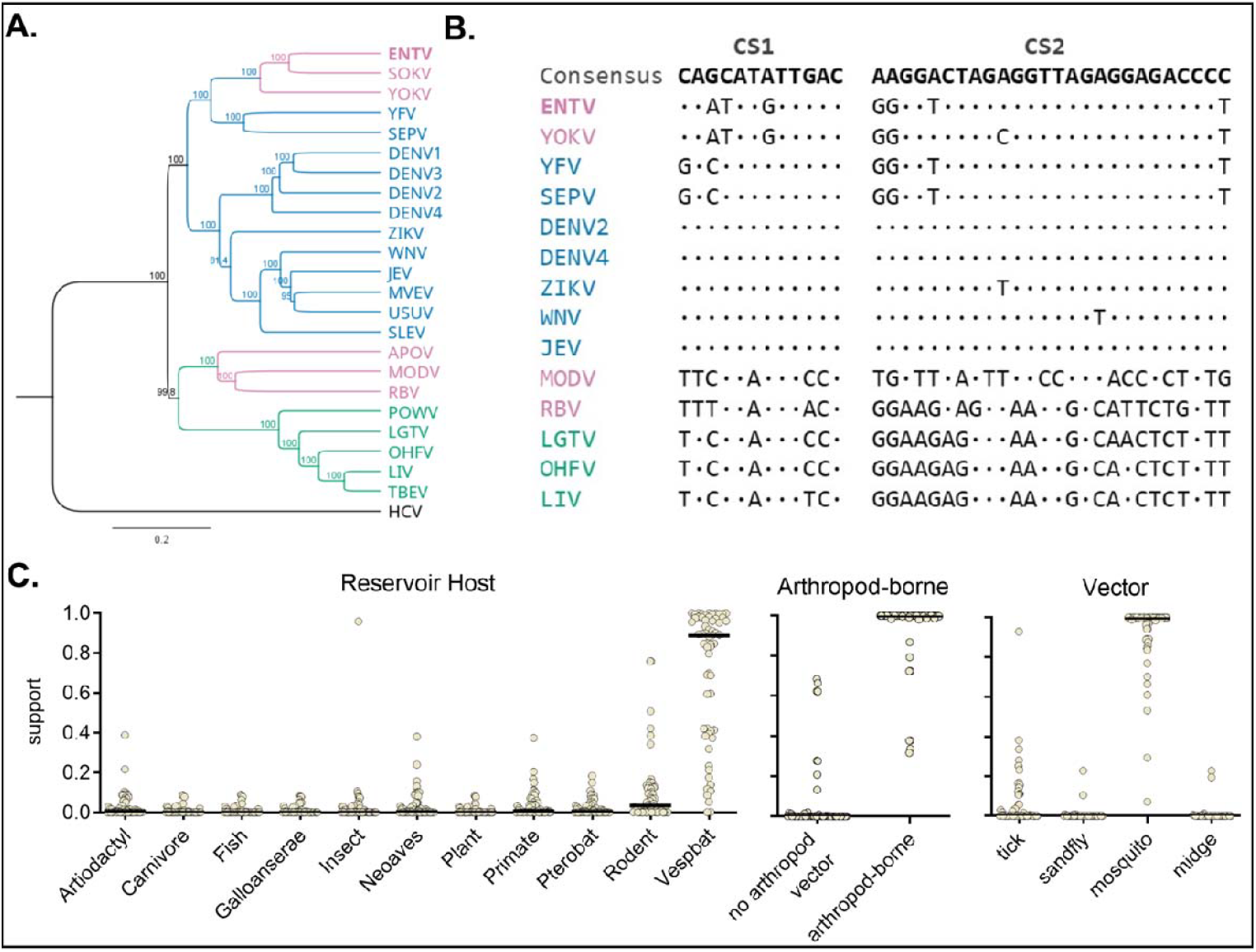
ENTV has characteristics consistent with mosquito-borne flaviviruses. **A)** Various mosquito-borne, tick-borne, and no-known vector flaviviruses nucleotide sequences were aligned with ENTV by the MUSCLE algorithm. The phylogenetic tree, rooted with hepatitis C virus as an outgroup, was constructed using the UPGMA method with 500 bootstraps in Geneious Prime. **B)** CS1 and CS2 alignment of flaviviruses generated by ClustalW alignment in MEGA12. Consensus from MBFs is shown at the top and each virus nucleotides are shown if differ from the consensus. The virus abbreviation and the accession numbers are listed in Supplementary Table 2. **C)** Host prediction support for reservoir species, presence of arthropod vector, and arthropod vector species using Viral Host Predictor by CVR Bioinformatics (https://bioinformatics.cvr.ac.uk/software/viral-host-predictor/)^23^. The bars on the graph represent the median.

### Experimental infection of mosquitoes

To determine the ability of mosquitoes to become infected with ENTV, *Ae. aegypti, Ae. albopictus*, and *Cx. quinquefasciatus*, were fed an infectious bloodmeal containing 1.1×10^6^ pfu/mL ENTV. At 7 days post-infection (dpi), whole bodies were collected and tested for viral RNA (vRNA) by qRT-PCR, and infectious virus by CPE assay. Although some mosquitoes (<25%) had vRNA levels above the background, all were negative for infectious virus (**Figure 4A, Table 2**). Next, we determined if ENTV can replicate in mosquitoes when injected intrathoracically (IT), bypassing the midgut infection barrier. *Ae. aegypti* and *Ae. albopictus* were IT inoculated with 2.3 × 10^2^ pfu ENTV, and whole bodies were tested for vRNA and infectious virus at 0, 7 and 14 dpi. Unlike bloodmeal exposure of ENTV, almost all (>95%) IT injected mosquitoes were vRNA positive for ENTV at 7 and 14 dpi, with most vRNA-positive mosquitoes also positive for CPE, confirming the presence of infectious ENTV (**Table 3**). vRNA increased over 14 days, by ∼100-fold and ∼400-fold in *Ae. aegypti* and *Ae. albopictus*, respectively (**Figure 4B**). To determine dissemination and transmission, vRNA and infectious virus was measured in carcass, legs and wings, and saliva from a separate set of mosquitoes at 14 dpi. ENTV RNA and infectious virus was detected in the carcass, and legs and wings of 100% of mosquitoes (**Figure 4C, Table 4**). While some saliva samples had low levels of ENTV RNA, none were positive for infectious virus (**Figure 4C, Table 4**).

**Table 1.**
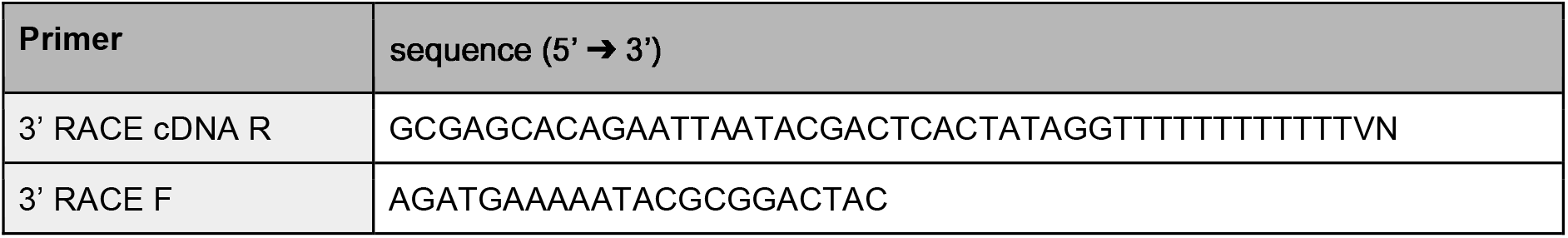
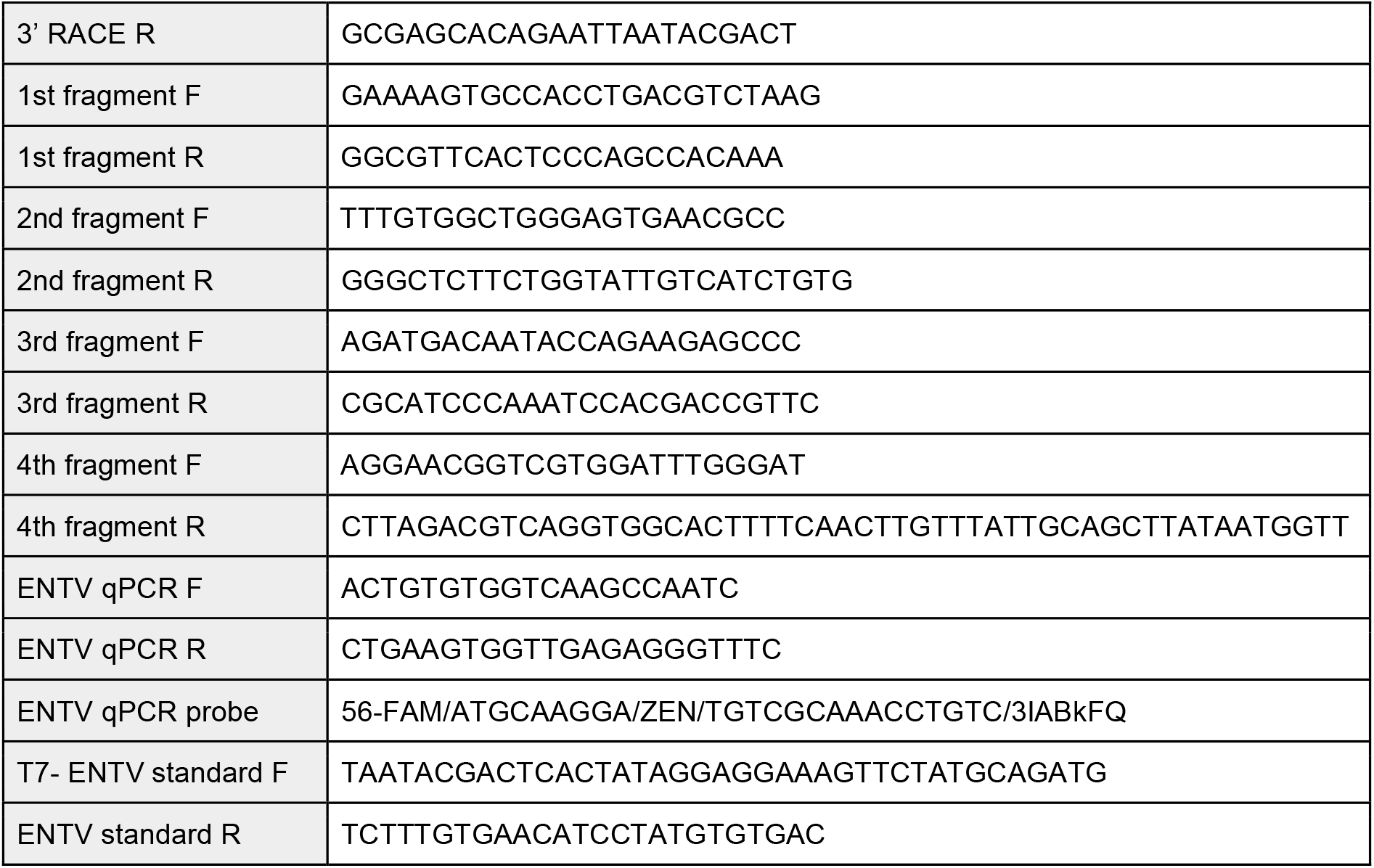
List of primers for 3’ RACE, qPCR, and reverse genetics PCR.

**Table 2.** ENTV in mosquito blood-feeding samples at 7dpi.

| Species | Positive by qPCR | Positive by CPE |
| --- | --- | --- |
| *Aedes aegypti* | 8/72 (11.1%) | 0/72 (0%) |
| *Aedes albopictus* | 16/72 (22.2%) | 0/72 (0%) |
| *Culex quinquefasciatus* | 5/72 (6.94%) | 0/72 (0%) |

**Table 3.** ENTV in IT-inoculated mosquito whole body.

| Species | days post inoculation | Positive by qPCR | Positive by CPE |
| --- | --- | --- | --- |
| *Aedes aegypti* | 0 | 6/6 (100%) | 6/6 (100%) |
|  | 7 | 47/48 (97.9%) | 45/48 (93.8%) |
|  | 14 | 48/48 (100%) | 47/48 (97.9%) |
| *Aedes albopictus* | 0 | 6/6 (100%) | 6/6 (100%) |
|  | 7 | 48/48 (100%) | 48/48 (100%) |
|  | 14 | 35/36 (97.2%) | 35/36 (97.2%) |

**Table 4.** ENTV in IT-inoculated mosquito samples at 14 dpi.

| Species | Part | Positive by qPCR | Positive by CPE |
| --- | --- | --- | --- |
| *Aedes aegypti* | carcass | 48/48 (100%) | 48/48 (100%) |
|  | legs and wings | 48/48 (100%) | 48/48 (100%) |
|  | saliva | 5/48 (10.4%) | 0/48 (0%) |
| *Aedes albopictus* | carcass | 48/48 (100%) | 48/48 (100%) |
|  | legs and wings | 48/48 (100%) | 47/48 (97.9%) |
|  | saliva | 19/48 (39.6%) | 0/48 (0%) |

**Figure 4.**
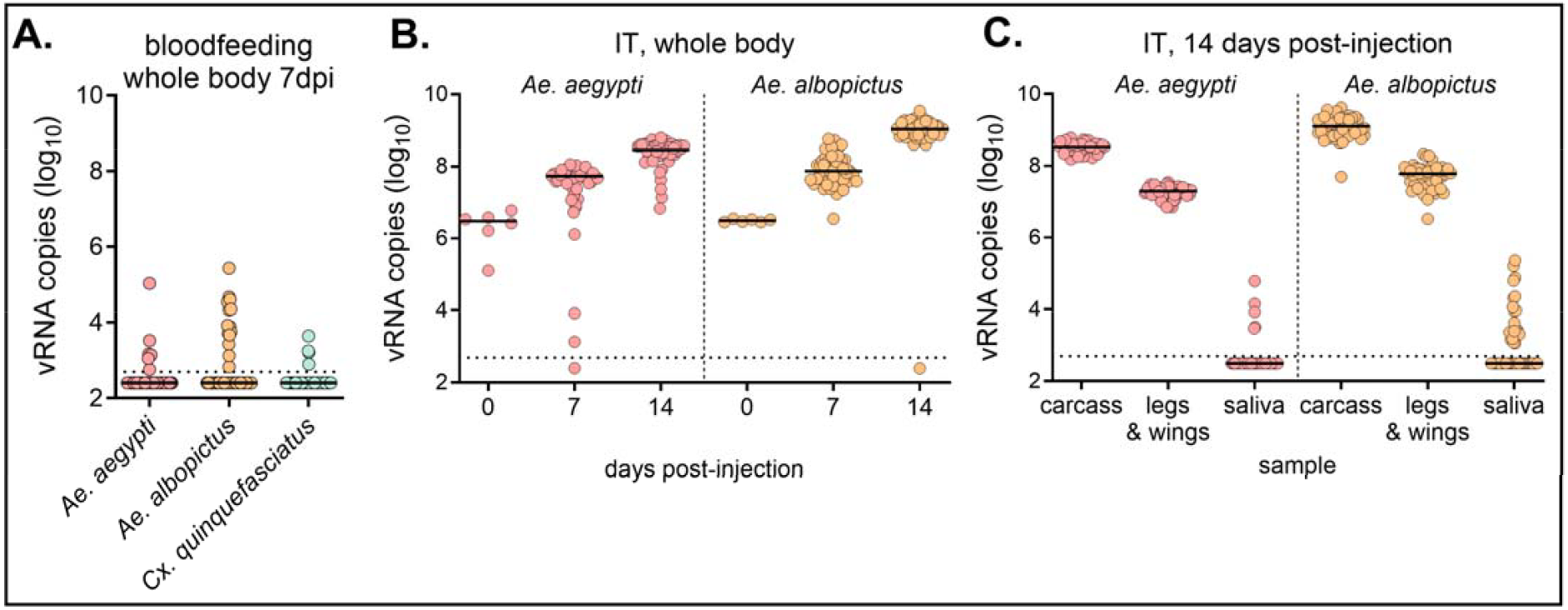
ENTV can replicate in mosquitoes but cannot overcome the midgut and salivary gland barriers. **A)** *Ae. aegypti, Ae. albopictus*, and *Cx. quinquefasciatus* were bloodfed with ENTV. At 7 dpi, mosquito whole bodies were homogenized and levels of ENTV RNA quantified (n=72, line shows median). **B, C)** *Ae. aegypti* and *Ae. albopictus* female were introthoracically inoculated with ENTV. **B)** Mosquitoes were collected 0, 7, and 14dpi, and ENTV RNA copy in the whole body was determined by qRT-PCR. **C)** Mosquitoes were salivated at 14dpi and ENTV RNA copy in the carcass, legs and wings, and saliva was determined by qRT-PCR. The dotted line shows the limit of detection.

## Discussion

Whether there is an arthropod host for ENTV and other NKV flaviviruses within the clade is intriguing, given their close phylogenetic relationships to known MBFs.^2,8,9,24^ Despite the interest in the host range of NKV flaviviruses, limited information is available on ENTV compatibility with cell lines^2,7^ or transmissibility in mosquito vectors.^6^ In this study, we developed an ENTV infectious clone and determined ENTV replication kinetics in various vertebrate and invertebrate cell lines. The second part of this study determined ENTV replication and biological barriers of infection in experimentally infected mosquitoes.

After failing to clone ENTV in a single plasmid, we developed a four-plasmid reverse genetics system. Increasing the number of fragments can affect the ligation efficiency in Gibson Assembly reactions; however, the recovery of the infectious clone was efficient and consistent across replicates, demonstrating the robustness of our system. Many use reverse genetics to generate attenuated live vaccines with multiple mutations.^25–27^ This process often entails multiple rounds of molecular cloning to generate single and combinatorial (e.g., double or triple) mutants, particularly in non-segmented viruses such as flaviviruses. This multiple plasmid system is advantageous in such applications because combinational mutants can be generated without additional cloning simply by substituting different mutation-containing fragments in the Gibson Assembly reaction, as long as the mutations are present on separate plasmids.

Consistent with previous reports^2,8,9^, phylogenetic analysis of the complete ENTV genome (including the newly sequenced UTR) suggested ENTV is closely related to MBFs. MBFs harbor CS1 and CS2 in the 3’ UTR^8,9,28^. The previously reported ENTV sequence contained CS2^2,8^ but terminated upstream of the expected CS1. This study determined the sequence for the rest of 3’ UTR, and confirmed the presence of the CS1 identical to that of YOKV.^24^ This finding further supports the close phylogenetic relationship between NKV flaviviruses in the ENTV clade and MBFs. Bioinformatics analysis, which considers genome sequence biases and phylogenetic relations with viruses with known hosts, suggested a mosquito host for ENTV. This finding is consistent with previously reported dinucleotide bias analysis^8^ and supports the growing speculation that ENTV and other NKV flaviviruses in the MBF clade have an unknown mosquito host.^8,29,30^

In our study, ENTV replicated well in multiple *Aedes* mosquito cell lines without requiring adaptation, reaching a titer ∼10^2^ higher than previously reported.^7^ ENTV also replicated in IT-inoculated mosquitoes, indicating that ENTV’s compatibility with mosquito cells is not limited to cell culture systems, which are often heavily adapted to laboratory conditions, and can be biased by its clonal and often immunodeficient population.^31^ Despite its molecular compatibility with mosquito cells, ENTV could not infect mosquitoes through blood feeding, a natural route of arbovirus infection. Furthermore, even when the midgut infection/escape barrier was artificially bypassed by IT-inoculation, ENTV failed to disseminate into saliva. For an arbovirus to be transmitted by mosquitoes to the next vertebrate host, it must successfully infect the mosquito midgut, escape to the haemocoel, reach and infect the salivary gland, and ultimately be released into saliva. A previous study demonstrated that *Aedes aegypti* is unable to transmit ENTV to mice via blood feeding, suggesting that transmission is blocked at either the midgut or salivary gland level.^26^ Our results from both blood feeding and IT inoculation indicate that ENTV is unable to overcome barriers at both the midgut and salivary gland stages. Therefore, we conclude that these mosquitoes (*Ae. aegypti* and *Ae. albopictus*) are not likely natural vectors of ENTV.

ENTV has only been isolated twice, both times from bats, and little is known about its ecology. Our study demonstrated efficient ENTV replication in *Aedes* mosquito cells; however, there is no convincing evidence to suggest vector transmission of ENTV. In Uganda, where ENTV was isolated, ∼200 mosquito species are reported, including 40 in the genus *Aedes* alone.^5,32,33^ Some MBFs have been isolated from diverse mosquito species in Africa, including non-Aedes mosquitoes from the genera *Coquillettidia* and *Mansonia*.^34–36^ Future work should evaluate if any of these mosquitoes can transmit ENTV. Additionally, our findings do not rule out the possibility of direct bat-to-bat transmission of ENTV, which, to the best of our knowledge, has not been previously investigated. Experimental studies exploring this potential mode of transmission would be valuable.

Finally, ENTV exhibited clear incompatibility with certain mosquito/tick cell lines, which are permissive to other flaviviruses.^37,38^ Many studies have used chimeric viruses to identify the molecular determinants of host block, which can be the incompatibility of structural proteins (block of binding and entry), nonstructural proteins (block post-entry, including replication and immune evasion), or interactions with the untranslated region.^12,29,39^ Since the host range is of particular interest in NKV flaviviruses, including ENTV, it would be interesting to use the reverse genetics system to study viral determinants of host specificity or adaptation to different hosts.

## Materials and Methods

### Virus

ENTV strain UG125 (GenBank accession KP233893.1) was initially isolated from a little free-tailed bat (*Mops pumilus*)^4^. The isolate was passaged twice in Vero cells. The virus used in this study was further passaged once in Vero and once in BHK cells, and contained a synonymous mutation A6,941G relative to the published sequence as determined by Nanopore sequencing on infected cells and the whole plasmid sequencing during the four-plasmid system construction.

### Cells

AJ6 is a primary cell line from a Jamaican fruit bat, *Artibeus jamaicensis*, and is a gift from Dr. Tony Schountz at Colorado State University. Mammalian cell lines (BHK21, Vero E6, Huh7, and AJ6) were propagated at 37°C in Dulbecco’s Modified Eagle’s Medium (DMEM). The DF-1 chicken fibroblast cell line was maintained in DMEM at 39°C. C6/36 (*Aedes albopictus*) cells were maintained at 28°C in DMEM supplemented with non-essential amino acids. Aag2 (*Aedes aegypti*) and CT (*Culex tarsalis*) cells were maintained at 28°C in Schneider’s Drosophila Medium. ISE6 (*Ixodes scapularis*) cells were cultured at 34°C in L15B-300 media supplemented with 5% triphosphate broth and 0.1% lipoprotein. All cell culture media was supplemented with 10% FBS, and except for ISE6, further supplemented with 100U/mL penicillin and 100µg/mL streptomycin. All cells, except for Aag2 and CT, were maintained with 5% CO_2_ with the addition of 25 mM HEPES (pH = 7.2) to the media.

### 3’ Rapid Amplification of cDNA Ends (RACE)

RNA was extracted from virus stock using TRIzol-LS (Invitrogen) following the manufacturer’s protocol. 3’ untranslated region (3’ UTR) of extracted RNA was polyadenylated with *E. coli* polyA polymerase (New England Biolabs) following the manufacturer’s protocol. Polyadenylated RNA was reverse transcribed with oligo dT primer with adapter sequence (**Table 1**) using Induro® Reverse Transcriptase (New England Biolabs) at 65°C for 2 hours. 3′ RACE was conducted using a forward primer targeting a known sequence in the 3′ UTR and a reverse primer designed on the adapter sequence (**Table 1**). PCR amplification was performed with Q5® High-Fidelity DNA Polymerase (New England Biolabs) under the following thermal cycling conditions: initial denaturation at 98°C for 30 seconds; 35 cycles of denaturation at 98°C for 10 seconds, annealing at 62°C for 30 seconds, and extension at 72°C for 30 seconds; followed by a final extension at 72°C for 2 minutes. The PCR product was purified using sparQ PureMag Beads (QuantaBio) and was Sanger sequenced by Azenta Life Sciences.

### Construction of the Four-Plasmid System

The four plasmids were cloned from viral cDNA or subcloned from a plasmid containing the partial ENTV genome in a CMV promoter-HDV ribozyme cassette. To develop the original single plasmid infectious clone, viral RNA was extracted from the virus stock using Trizol-LS (Invitrogen). cDNA was generated from random hexamers using Induro® Reverse Transcriptase (New England Biolabs) at 65°C for 2 hours. The ENTV genome was PCR-amplified in five overlapping fragments and joined by overlap PCR using Q5® High-Fidelity DNA Polymerase (New England Biolabs). The entire ENTV genome was inserted in a backbone containing a CMV promoter and HDV ribozyme^15^ by Gibson assembly and transfected into *E. coli (Stbl3*, ThermoFisher). This clone, containing a nonsense mutation and a large deletion, was used as a template to subclone partial genome fragments into plasmids 1 (pBG878), 2 (pBG879), and 4 (pBG881). The deleted genomic region was PCR amplified from viral cDNA and cloned into plasmid 3 (pBG880) (primers listed in **Supplementary Table 1**). Each plasmid was amplified in *E. coli*, harvested, and the whole plasmid sequence was verified by Plasmidsaurus.

### Generation of Recombinant Virus

From each plasmid, the partial ENTV genome with ∼30bp overlap at both ends was PCR amplified with Q5 DNA polymerase. Equimolar product was pooled into a total of ∼500ng in 15µL and mixed with an equal volume of NEBuilder® HiFi DNA Assembly Master Mix (New England Biolabs). The reaction was incubated at 50°C for 150 minutes and subsequently transfected into 80% confluent BHK cells in a 12-well plate using Lipofectamine 2000. At 12 hours post-transfection, cell culture media was replaced with 2% FBS DMEM, and infectious virus in the supernatant was collected every 24 hours. We named our infectious clone ENTV_IC and deposited the sequence in GenBank under accession number PQ720541.

### Growth Curve

Growth curves were generated for each cell line using three independent infections. Virus dilutions were prepared in 2% FBS media to achieve a multiplicity of infection (MOI) of 0.01. Sub-confluent cell monolayers were inoculated with the virus solution and incubated for 1 hour at the appropriate propagation temperature for each cell line. After inoculation, cells were washed five times with phosphate-buffered saline (PBS) containing 2% FBS and overlaid with 2% FBS media. Virus samples were collected at specified time points and stored at -80°C until titration. The infectious titer of each sample was determined using an agarose plaque assay performed on BHK cells in a 12-well plate. Five days post-infection, plates were fixed and stained with 0.1% crystal violet for visualization.

### RNA extraction and qRT-PCR

RNA from mosquito samples was extracted using Mag-Bind® Viral DNA/RNA 96 Kit (Omega Bio-Tek) using the KingFisher Flex Magnetic Particle Processor (ThermoFisher). All qRT-PCR reactions were performed using the EXPRESS One-Step Superscript™ qRT-PCR Kit (Invitrogen), following the manufacturer’s instructions. Final concentrations of primers and probe were 50 nM and 17.5 nM, respectively. Thermal cycling conditions were as follows: reverse transcription at 50°C for 15 minutes, enzyme activation at 95°C for 3 minutes, followed by 40 cycles of amplification at 95°C for 5 seconds and 72°C for 30 seconds. For each viral target, an RNA standard (∼1,000 kb) was synthesized using T7 polymerase, and RNA copy numbers were calculated based on the corresponding standard curve. All qRT-PCR primers, probes, and primers used for RNA standard synthesis are listed in Table 1.

### Mosquitoes

We used three laboratory colonies: *Culex quinquefasciatus* (>50 passages, established from a colony maintained by WK Reisen collected in 1953 from California), *Ae. aegypti* Poza Rica (originally collected from Veracruz, Mexico in 2012^43^), and *Ae. albopictus* strain ATM NJ-95 (F56, initially collected in New Jersey, USA in 1995. Eggs were obtained from BEI / F39 Cat No. NR-48979). Mosquitoes were maintained at 26– 27°C (*Cx. quinquefasciatus*) or 28°C (*Aedes* spp.) with relative humidity of 70–80% and given raisins and water. Larvae were raised on powdered fish food.

### Mosquito blood feeding

Seven to ten days after emergence, mosquitoes were given warmed virus-containing bloodmeal using a glass water-jacketed membrane feeding apparatus sealed with hog’s gut. Virus was freshly produced by passaging ENTV stock on BHK cells (MOI = 0.01) and harvesting 2 days post-infection. Bloodmeal consisted of a 1:1 mixture of virus (final concentration of 1.1 × 10^6^ pfu/mL) and defibrinated calf’s blood and was supplemented with 1 mM ATP. Mosquitoes were allowed to feed for approximately 30 minutes. Fully engorged females were sorted and were maintained at 27°C with relative humidity at 70–80% with sucrose and water. Mosquitoes were harvested at the noted time points post-blood-feeding, and the whole body was homogenized in mosquito diluent (PBS supplemented with 20% FBS, 2.5µg/mL amphotericin B, 50µg/ml gentamicin) with a metal bead at 24Hz per second for 1 minute. The resultant homogenate was plated onto BHK cells, and infection was confirmed by observing cytopathic effect, followed by qRT-PCR as described above.

### Intrathoracic inoculation

Five to six days post-emergence, adult females were intrathoracically inoculated with 2.3 × 10^2^ pfu of ENTV in a volume of 69 nL using a Nanoject II (Drummond Scientific Company, Broomall, PA, USA).^40^ All injections were performed under a dissecting microscope using glass needles prepared using a vertical pipette puller (P-30, Sutter Instrument Co., Novato, CA, USA). Mosquitoes were maintained, and samples were harvested and processed as described above.

### Mosquito salivation/dissection

At 14 days post-IT inoculation, mosquito legs and wings were removed with forceps under cold anesthesia. The mosquito was then forced to salivate in a glass capillary containing mineral oil for 1 hour at 27°C with relative humidity at 70–80%. Salivation was confirmed by observing the capillary glass under a microscope, and saliva was ejected into 250μL of 20% FBS DMEM using a dropper pipette. Legs and wings, and post-salivation carcasses were processed for homogenization as described in the mosquito blood-feeding section. All samples were tested for vRNA and infectious virus as described above.

## Acknowledgements

We would like to thank the US Centers for Disease Control Division of Vector-borne Diseases Arbovirus Reference Collection for the ENTV isolate, Dr. Tony Schountz (Colorado State University) for supplying AJ6 cells, and Dr. Emma Harris (Colorado State University) for ISE6 cell culture assistance.

## Financial Support

This research was supported by NIH under grant numbers R01 AI166050 and R01 AI067380. M.F. was also supported by the Takenaka Scholarship Foundation. ENG was supported by National Science Foundation DBI 2021909, 2213854, and 2515340.

## Disclosures

The authors declare no conflicts of interest.

## Author contributions

MF: conceptualization, investigation, methodology, writing – original draft

ENG: investigation, writing – review & editing

ISV: investigation

BME: investigation

LEM: investigation

GDE: supervision, project administration, funding acquisition

BJG: conceptualization, supervision, project administration, funding acquisition

## Notes

### Competing Interest Statement

The authors have declared no competing interest.

