## Supplemental Data for "Development of ENTV reverse genetics system and phenotypic evaluation of rescued virus reveals host-specific replication patterns in mosquitoes"

### Slide 1
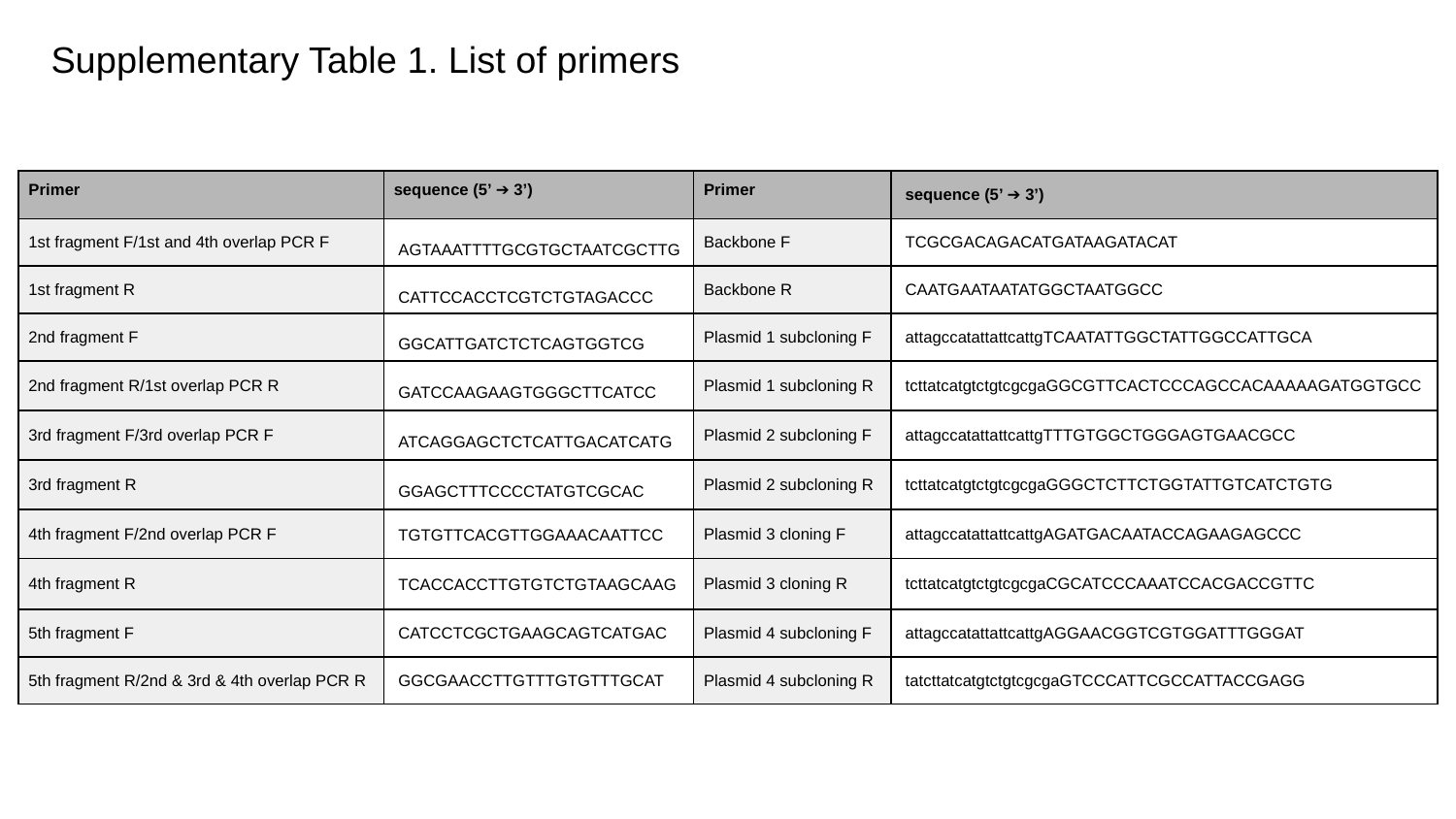

#### Supplementary Table 1. List of primers
| Primer | sequence (5’ ➔ 3’) | Primer | sequence (5’ ➔ 3’) |
| --- | --- | --- | --- |
| 1st fragment F/1st and 4th overlap PCR F | AGTAAATTTTGCGTGCTAATCGCTTG | Backbone F | TCGCGACAGACATGATAAGATACAT |
| 1st fragment R | CATTCCACCTCGTCTGTAGACCC | Backbone R | CAATGAATAATATGGCTAATGGCC |
| 2nd fragment F | GGCATTGATCTCTCAGTGGTCG | Plasmid 1 subcloning F | attagccatattattcattgTCAATATTGGCTATTGGCCATTGCA |
| 2nd fragment R/1st overlap PCR R | GATCCAAGAAGTGGGCTTCATCC | Plasmid 1 subcloning R | tcttatcatgtctgtcgcgaGGCGTTCACTCCCAGCCACAAAAAGATGGTGCC |
| 3rd fragment F/3rd overlap PCR F | ATCAGGAGCTCTCATTGACATCATG | Plasmid 2 subcloning F | attagccatattattcattgTTTGTGGCTGGGAGTGAACGCC |
| 3rd fragment R | GGAGCTTTCCCCTATGTCGCAC | Plasmid 2 subcloning R | tcttatcatgtctgtcgcgaGGGCTCTTCTGGTATTGTCATCTGTG |
| 4th fragment F/2nd overlap PCR F | TGTGTTCACGTTGGAAACAATTCC | Plasmid 3 cloning F | attagccatattattcattgAGATGACAATACCAGAAGAGCCC |
| 4th fragment R | TCACCACCTTGTGTCTGTAAGCAAG | Plasmid 3 cloning R | tcttatcatgtctgtcgcgaCGCATCCCAAATCCACGACCGTTC |
| 5th fragment F | CATCCTCGCTGAAGCAGTCATGAC | Plasmid 4 subcloning F | attagccatattattcattgAGGAACGGTCGTGGATTTGGGAT |
| 5th fragment R/2nd & 3rd & 4th overlap PCR R | GGCGAACCTTGTTTGTGTTTGCAT | Plasmid 4 subcloning R | tatcttatcatgtctgtcgcgaGTCCCATTCGCCATTACCGAGG |

### Slide 2
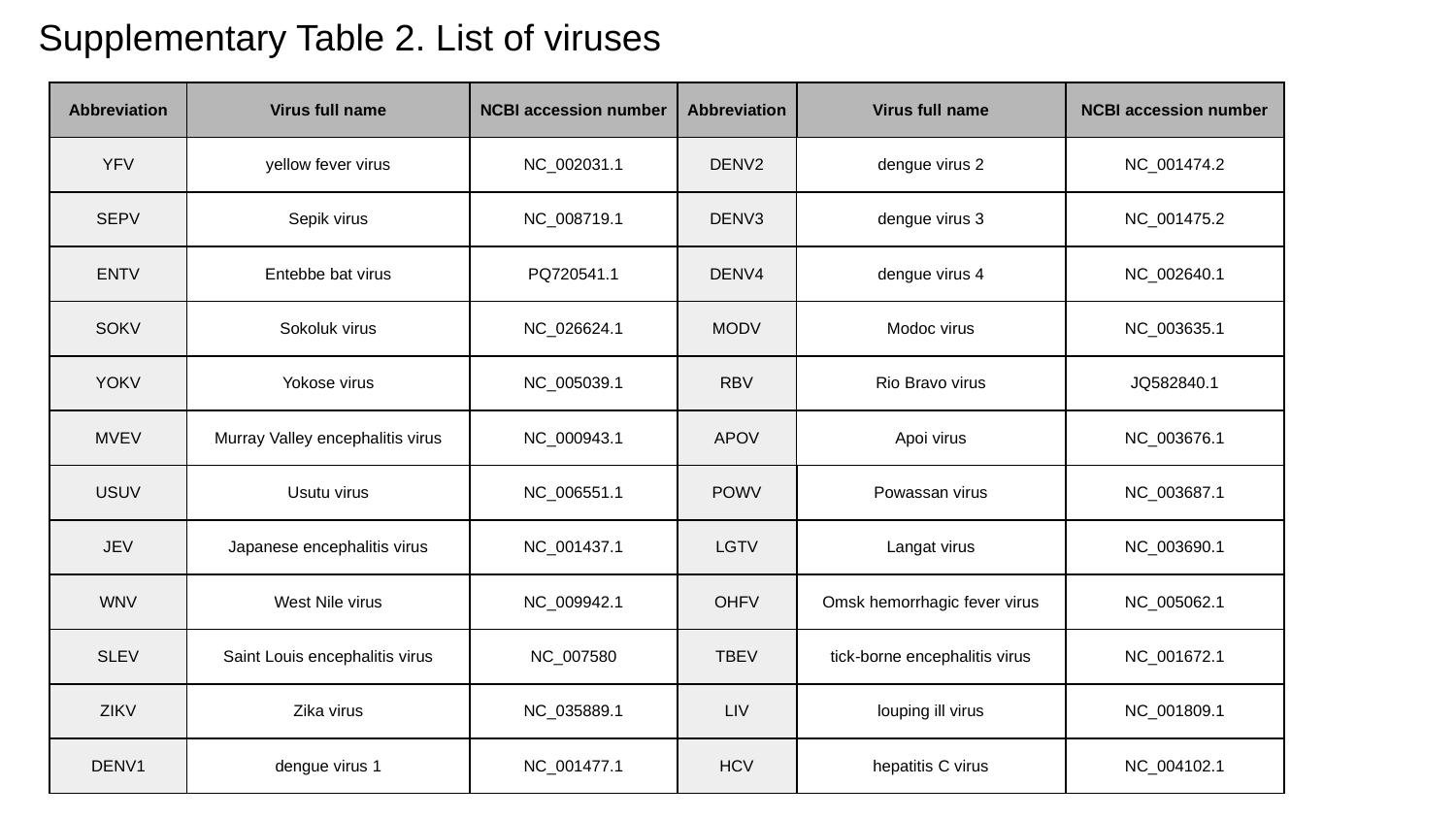

#### Supplementary Table 2. List of viruses
| Abbreviation | Virus full name | NCBI accession number | Abbreviation | Virus full name | NCBI accession number |
| --- | --- | --- | --- | --- | --- |
| YFV | yellow fever virus | NC\_002031.1 | DENV2 | dengue virus 2 | NC\_001474.2 |
| SEPV | Sepik virus | NC\_008719.1 | DENV3 | dengue virus 3 | NC\_001475.2 |
| ENTV | Entebbe bat virus | PQ720541.1 | DENV4 | dengue virus 4 | NC\_002640.1 |
| SOKV | Sokoluk virus | NC\_026624.1 | MODV | Modoc virus | NC\_003635.1 |
| YOKV | Yokose virus | NC\_005039.1 | RBV | Rio Bravo virus | JQ582840.1 |
| MVEV | Murray Valley encephalitis virus | NC\_000943.1 | APOV | Apoi virus | NC\_003676.1 |
| USUV | Usutu virus | NC\_006551.1 | POWV | Powassan virus | NC\_003687.1 |
| JEV | Japanese encephalitis virus | NC\_001437.1 | LGTV | Langat virus | NC\_003690.1 |
| WNV | West Nile virus | NC\_009942.1 | OHFV | Omsk hemorrhagic fever virus | NC\_005062.1 |
| SLEV | Saint Louis encephalitis virus | NC\_007580 | TBEV | tick-borne encephalitis virus | NC\_001672.1 |
| ZIKV | Zika virus | NC\_035889.1 | LIV | louping ill virus | NC\_001809.1 |
| DENV1 | dengue virus 1 | NC\_001477.1 | HCV | hepatitis C virus | NC\_004102.1 |

### Slide 3
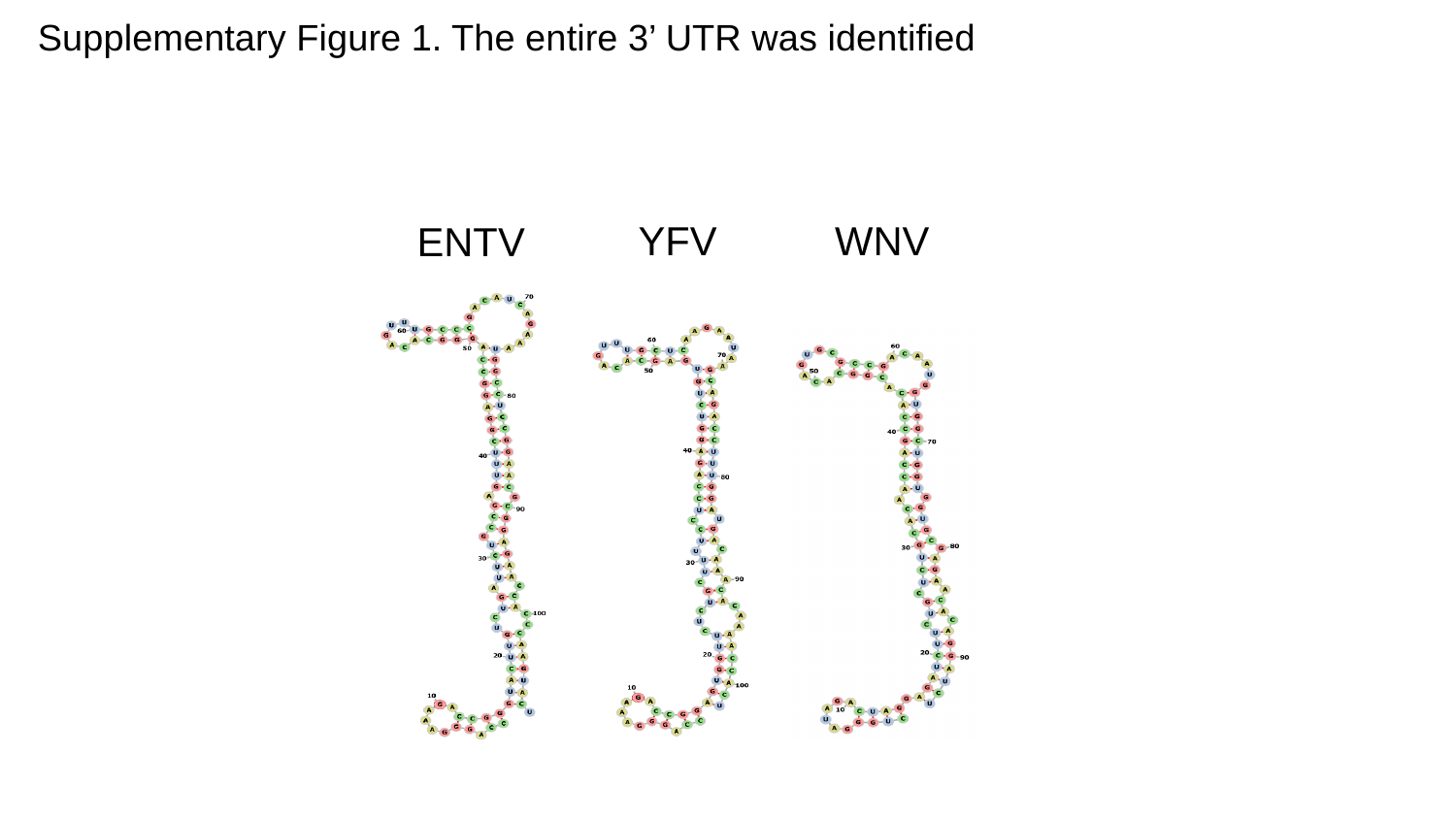

#### Supplementary Figure 1. The entire 3’ UTR was identified
YFV
ENTV
WNV

### Slide 4
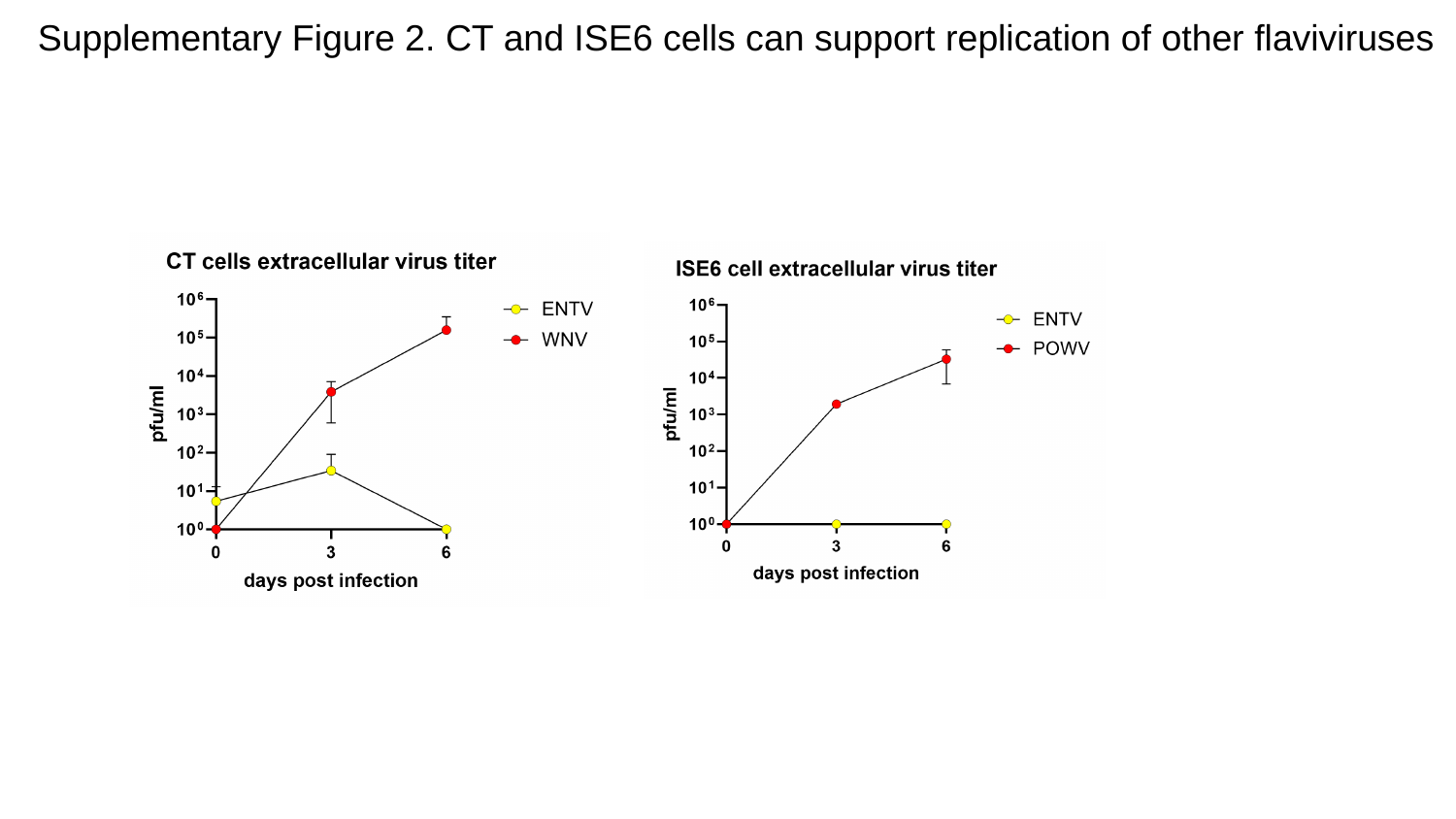

#### Supplementary Figure 2. CT and ISE6 cells can support replication of other flaviviruses

### Slide 5
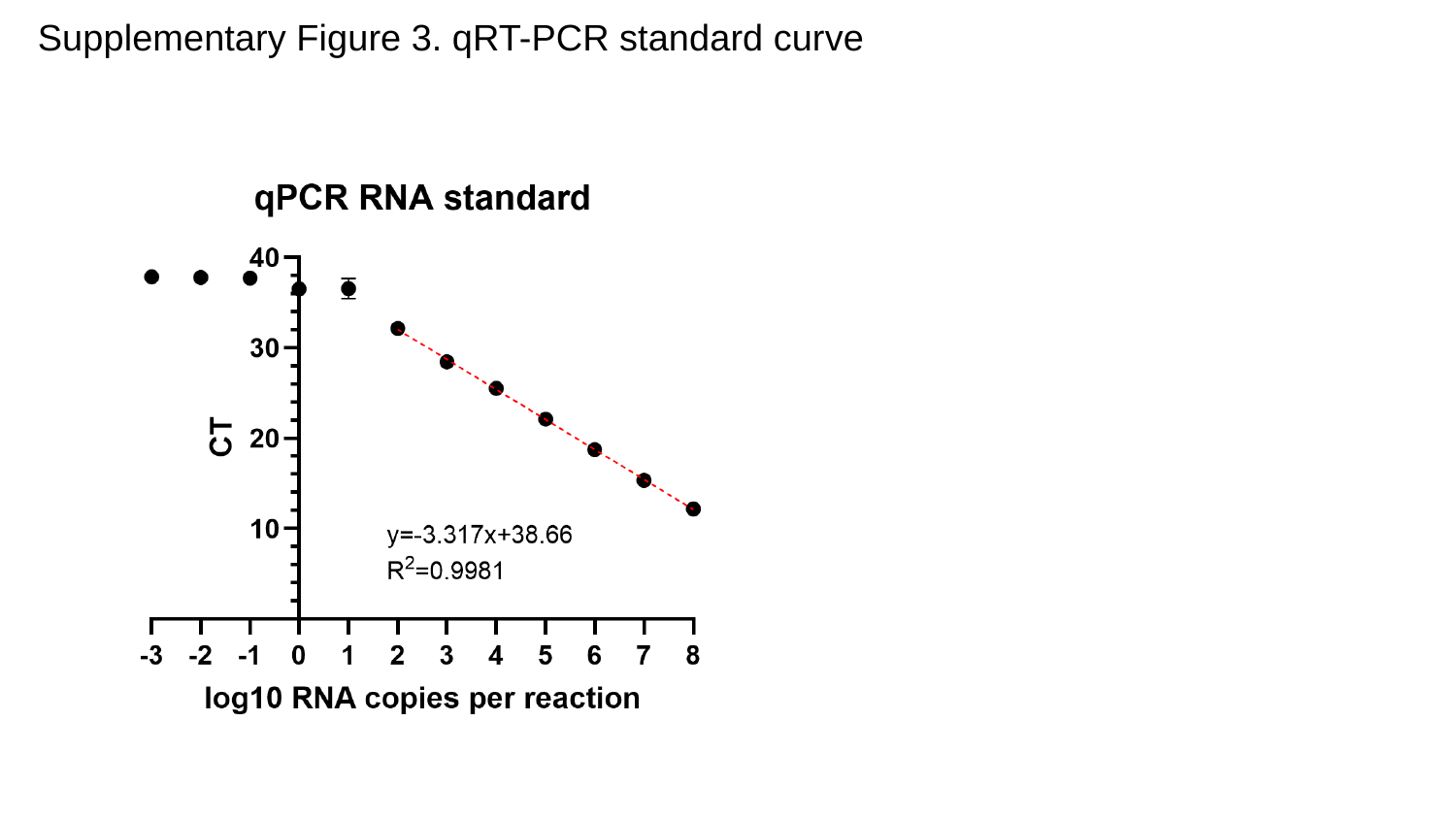

#### Supplementary Figure 3. qRT-PCR standard curve
